# Genome-wide analysis reveals genetic diversity, linkage disequilibrium, and selection for milk traits in Chinese buffalo breeds

**DOI:** 10.1101/701045

**Authors:** Xing-Rong Lu, An-Qin Duan, Sha-Sha Liang, Xiao-Ya Ma, Xian-Wei Liang, Ting-Xian Deng

## Abstract

Water buffalo holds the tremendous potential of milk and meat that widespread throughout central and southern China. However, characterization of the population genetics of Chinese buffalo is poorly understood. Using Axiom^®^ buffalo genotyping array, we performed the genetic diversity, linkage disequilibrium (LD) pattern and signature of selection in the 176 Chinese buffaloes from thirteen breeds. A total of 35,547 SNPs passed quality control and were used for further analyses. Population genetic analysis revealed a clear separation between the swamp and river types. Ten Chinese indigenous breeds clustered into the swamp group, Murrah and Nili-Ravi breeds were the river group, and the crossbred breed was closer to the river group. Genetic diversity analysis showed that the swamp group had a lower average expected heterozygosities compared to the river group. LD decay distance was much shorter in the swamp group compared with the river group with 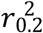 value of approximately 50 Kb. Analysis of runs of homozygosity indicated that extensive remote and recent inbreeding activity was respectively found within swamp and river groups. Moreover, a total of 12 genomic regions under selection were detected between river and swamp groups. Further, 12 QTL regions were found associated with buffalo milk production traits. Some candidate genes within these QTLs were predicted to be involved in the cell structure and function, suggesting that these genes might play vital roles in the buffalo milk performance. Our data contribute to our understanding of the characterization of population genetics in Chinese buffaloes, which in turn may be utilized in buffalo breeding programs.

**Author Summary:** Identifying the causal genes or markers associated with important economic traits in livestock is critical to increasing the production level on the species. However, current understanding of the genetic basis for milk production traits in buffalo is limited. Here, we confirmed the divergent evolution, distinct population structure, and LD extent among Chinese buffalo breeds. We also identified 12 QTL regions associated with milk production traits in buffaloes using the selective sweeps and haplotype analysis. Further, a total of 7 genes involved in the cell structure and function were predicted within the identified QTLs. These findings suggested that these genes can serve as the candidate genes associated with buffalo milk production, which hold a vital role in the milk trait improvement of dairy buffalo industry.

## Introduction

Water buffalo (*Bubalus bubalis*), primarily raised for milk, meat and draught power, is an essential part of the agricultural economy of many countries around the world. Broadly, buffalo mainly consists of two types: the river (*B. bubalis bubalis*; n=50) and swamp (*B. bubalis carabensis*; n=48), which are primarily distinguished based on their distinct morphology and chromosomal karyotypes [1]. In China, the indigenous buffaloes were mainly the swamp type, with the largest number of swamp population across the world. To date, Chinese swamp buffaloes have been classed into 24 local types based mainly on the regional distribution [2]. To improve the milk and meat performance of swamp buffalo, crossbred buffalo by hybridizing the river-type bulls with swamp-type cows has been practiced in many Asian countries, and they are fertile, although the hybrid may have a lower reproductive value [3, 4]. The cross-breeding programs in China began in the 1950s by introducing the exotic river breeds such as Murrah and Nili-Ravi, which has resulted in the different crossbred breeds including the Murrah × local buffalo, Nili-Ravi × indigenous buffalo, and Murrah × Nili-Ravi × indigenous buffalo. These animals have higher milk production (~1,700 kg per lactation) than that of swamp buffalo (~500-800 kg per lactation) after multiple cross breeding for several decades, which is still lower than that of the river buffalo breeds (~2,200 kg/lactation) [5]. Therefore, it is of great interest to genetically dissect the molecular basis of buffalo complex traits such as milk and meat, which contribute to the development of dairy buffalo industry.

Knowledge of genetic diversity within and among breeds and populations is crucial for their domestication, conservation, and management. In the past few decades, two independent domestication events of the river and swamp buffaloes were confirmed using the Y-chromosomal [6] and mitochondrial DNA [7] technologies. River buffalo was domesticated in the western region of the Indian subcontinent [8], while swamp buffalo was domesticated in the border between Indochina and Southwestern China occurred 4000 years ago [9, 10]. More importantly, a robust geographic differentiation was found within the swamp buffaloes [9], and the Southwestern buffalo populations in China had the highest genetic diversity compared to the other domestication centers (Central and Southwestern China) [11]. These domestication events, including the natural and artificial selection, help to shape and stabilize the buffalo breed characteristics. For example, river buffalo is well known to be a breed that is mainly used for milk and meat production, while swamp buffalo is essentially a draught animal with lower milk production. With the acceleration of urbanization, however, swamp buffalo population in China shows a rapid decline trend in recent years, especially some endangered swamp breeds emerged. It is well known that a drop in population size may generally cause the decline of genetic diversity by genetic drift and inbreeding [12]. Because of genetic diversity underpins population resilience and persistence, it is of considerable importance to investigate the genetic diversity of swamp buffalo breed in China. These results will play a vital role in the buffalo genetic improvement, particularly in its breeding programs.

In the last decade, genome/transcriptome-wide molecular markers were identified using high-throughput sequencing [13–15], which can be utilized for animal breeding and genetic studies. Deng et al. [16] identified 17,401 simple sequence repeats (SSRs) in swamp buffalo that could be used as potential molecular markers by the Illumina paired-end technology that could help spearhead molecular genetics research studies on this species. Single nucleotide polymorphism (SNP) is another critical marker that can be used in genetic diversity studies or genetic mapping. Various genotyping techniques have also been developed for SNP discovery, ranging from the low-throughput allele-specific PCR [17] to high-throughput methods of genotyping hundreds of thousands of SNPs in parallel [18]. To date, high-throughput SNP genotyping has been extensively applied to various domestic animals [19–21] and plants [22]. The first SNP genotyping platform (Axiom® buffalo genotyping array; 90K Affymetrix) designed explicitly for river buffalo has recently been developed [23] and used for the genome-wide analysis study (GWAS) in different buffalo breeds [24–26]. This SNP90K array provides a viable choice for buffalo scientific research such as molecular breeding, complex traits, conservation, and biodiversity.

Several methods have been developed to detect signatures of recent selection in domesticated animals [27]. These methods mainly based on the linkage disequilibrium (LD), spectra of allele frequencies, and characteristics of haplotype structures in the studied populations [28]. Notably, identifying signatures of selection could provide insight into the genomic response to domestication and selection for production traits, which help in the design of more efficient selection schemes [29]. To date, numbers selective sweep regions associated with production traits has been identified in domesticated animals, such as milk production traits [30], reproductive traits [31], feed efficiency [32], elongation of the back [33], and lack of horns [34]. For the Azeri and Khuzestani buffalo breeds, Mokhber et al. [35] identified 13 selective sweep regions that are potentially related to economically important traits using the genome-wide SNP data. However, the selection footprints among the Chinese buffalo breeds remain unexplored. Therefore, we investigated thirteen buffalo breeds in South China region using the buffalo SNP90K array, aiming to explore the genetic diversity, LD extent, signature of selection and quantitative trait locus (QTL) among the studied breeds, which will benefit in the development of an SNP genotyping panel for swamp buffalo and promote the buffalo complex traits research and breeding program.

## Results

### SNP Characteristics

Statistic information on 176 buffaloes representing 13 breeds was summarized in **Table 1**. SNP information regarding the chromosomes, numbers, and length was listed in the **S1 Table**. A total of 35,547 SNPs was generated after filtering that covered 1309.75 Mb with an average distance of 36.84 Kb between adjacent SNPs. The average chromosomal length ranged from 21.97 Mb on Chr24 to 102.63 Mb on Chr1. The mean length of adjacent SNPs per chromosome ranged between 32.69 to 54.63 Kb on Chr19 and ChrX, respectively. The average LD was 0.21 across the buffalo genome.

**Table 1.** Genetic diversities for the analyzed buffalo populations in the South China region. N, Sample size after QC; No. of usable loci, number of loci with <5% of missing data; No. of polym. loci, number of loci that resulted polymorphic in a breed; Ho, observed heterozygosity He, expected heterozygosity; A_R_, allelic richness.

| Group | Breed | Code | N | No. of usable loci | No. of polym. loci | Ho(±SD) | He(±SD) | A<sub>R</sub> |
| --- | --- | --- | --- | --- | --- | --- | --- | --- |
| River |  |  | 43 | 35191 | 18081 | 0.429±0.11 | 0.436±0.08 | 1.994 |
|  | Murrah | MUR | 12 | 33250 | 16866 | 0.446±0.17 | 0.427±0.11 | 1.987 |
|  | Nili-Ravi | NIR | 11 | 34105 | 17008 | 0.414±0.18 | 0.401±0.12 | 1.969 |
|  | Crossbred | CRO | 20 | 34165 | 17573 | 0.438±0.14 | 0.433±0.09 | 1.999 |
| Swamp |  |  | 133 | 35544 | 13505 | 0.167±0.18 | 0.174±0.19 | 1.740 |
|  | Hunan | HUN | 15 | 35097 | 8073 | 0.284±0.19 | 0.284±0.17 | 1.452 |
|  | Ensi | ENS | 15 | 34978 | 6532 | 0.321±0.18 | 0.328±0.15 | 1.366 |
|  | Fuling | FUL | 15 | 35122 | 6483 | 0.323±0.18 | 0.330±0.15 | 1.361 |
|  | Yibin | YIB | 15 | 34963 | 6427 | 0.322±0.18 | 0.327±0.15 | 1.363 |
|  | Yangzhou | YAN | 14 | 31871 | 6622 | 0.311±0.19 | 0.306±0.16 | 1.426 |
|  | Dechang | DEC | 12 | 34611 | 10084 | 0.250±0.19 | 0.250±0.17 | 1.566 |
|  | Dehong | DEH | 12 | 34622 | 7688 | 0.309±0.18 | 0.315±0.16 | 1.438 |
|  | Fuzhong | FUZ | 12 | 33829 | 6401 | 0.329±0.18 | 0.332±0.15 | 1.376 |
|  | Xilin | XIL | 11 | 34590 | 6227 | 0.335±0.18 | 0.345±0.15 | 1.354 |
|  | Guizhou | GUI | 12 | 34665 | 6468 | 0.316±0.18 | 0.334±0.15 | 1.369 |

### Population analysis

Principal components analysis (PCA) showed a distinct separation between the river and swamp types (**Fig 1A**). The crossbred buffaloes were originated from the swamp- and river-type buffaloes and were concordantly located between them. Similar genetic relationship among analyzed breeds was also supported by phylogenetic analysis (**Fig 1B**) and population structure analysis (**Fig 1C**). As showed in **Fig 1C**, K=2 represented the most appropriate population number for the present dataset, indicating that there was an apparent differential distribution between river and swamp types. This suggested that the studied buffalo breeds can be divided into two groups (river and swamp) for further analysis.

**Fig 1.**
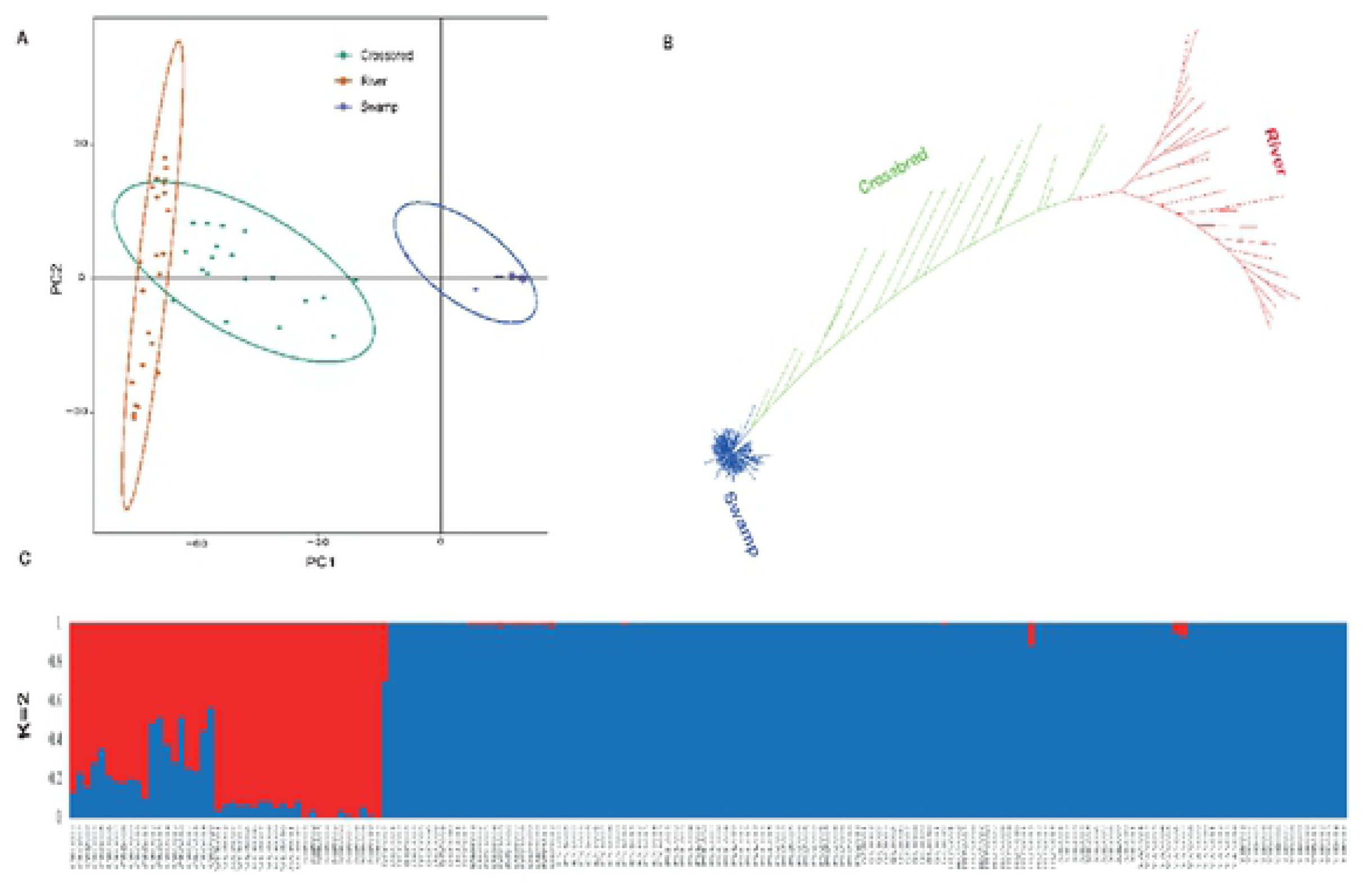
Population analysis of 13 buffalo breeds in South China. (A) PCA plots display the individuals’ relationship of 176 buffaloes. (B) Neighbor-joining representation of the pairwise Nei’s D genetic distances among populations. (C) Population structure of 176 buffaloes inferred by model-based clustering using ADMIXTURE.

### Genetic diversity

Distribution of 5 minor allele frequency (MAF) classes between river and swamp groups were presented in **Fig 2**. Compared to the river group, buffalo breeds in swamp group had the highest proportion of SNPs with lower MAF category (0, 0.1]. Buffalo breeds in river group had a relatively high proportion of SNPs with high MAF (mostly (0.2, 0.3], (0.3, 0.4], and (0.4, 0.5]).

**Fig 2.**
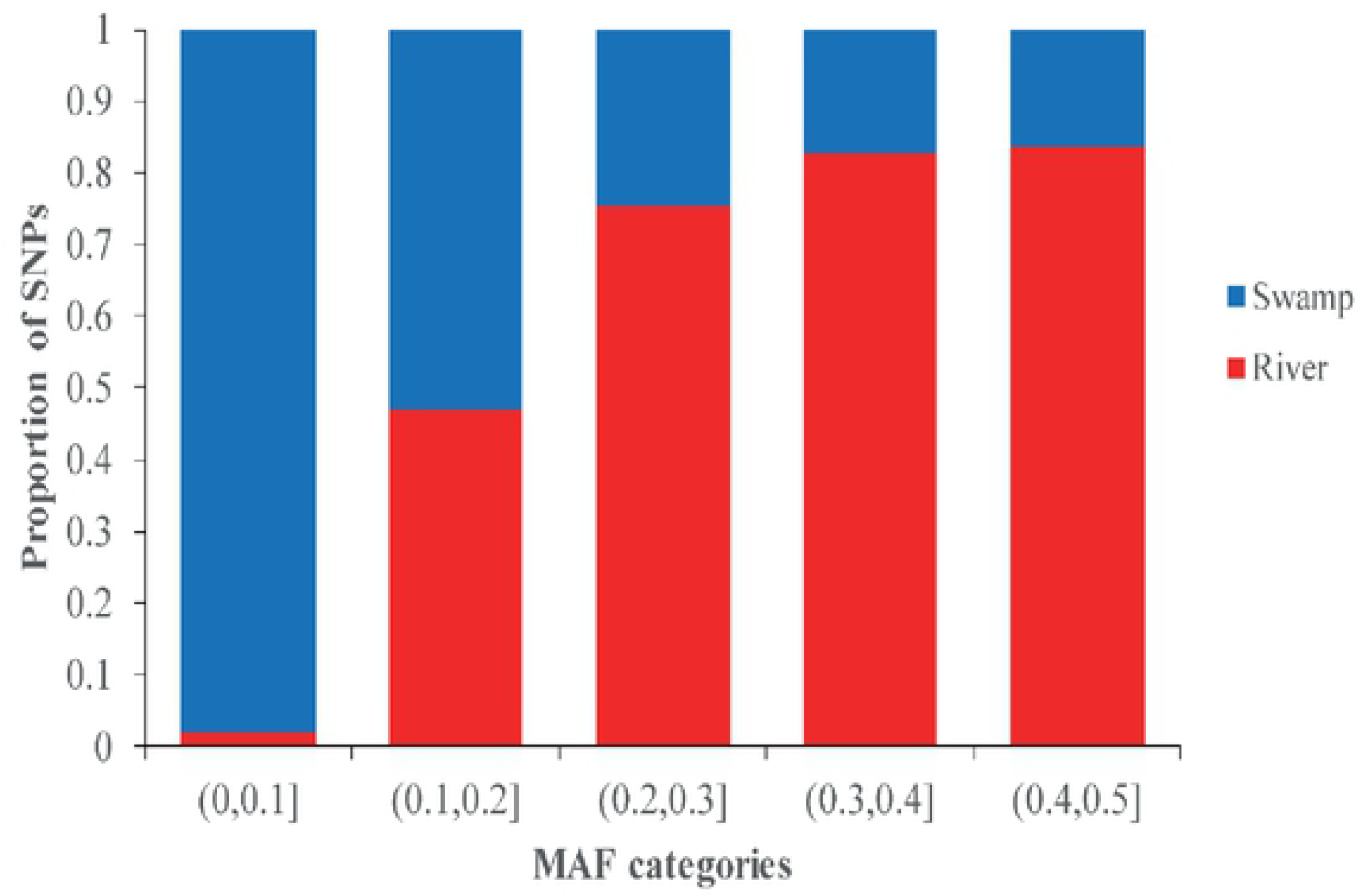
MAF Distribution for the river and swamp groups.

Genome-wide genetic diversity metrics within breeds were measured by the observed heterozygosity (H_O_), expected heterozygosity (He), and allelic richness (A_R_) (**Table 1**). Overall, buffaloes in river group displayed a comparably high level of polymorphism SNPs (50.87%) compared to the swamp group (37.99%). They had higher genetic diversity compared to the swamp group, as measured by the H_O_ (0.429 ± 0.11 *vs.* 0.167 ± 0.18) and He (0.436 ± 0.08 *vs.* 0.174 ± 0.19). The average heterozygosity estimates were highest for Murrah breed in the river group (H_O_: 0.446 ± 0.17; He: 0.427 ± 0.11) and lowest for DEC breed in swamp group (H_O_: 0.250 ± 0.19; He: 0.250 ± 0.17). The allelic richness for swamp group (A_R_ = 1.740) was lower than that of the river group (A_R_ = 1.994). Notably, the DEC breed in swamp group was observed to have the lowest A_R_ (A_R_ = 1.556), while the XIL breed revealed the lowest A_R_ (A_R_ = 1.354).

Population differentiation estimates showed that the pairwise *F*_ST_ values ranged from 0.0076 to 0.7535 across the studied breeds (**S2 Table**). For the swamp group, HUN was most closely related to FUZ (*F*_ST_ = 0.0087), GUI (*F*_ST_ = 0.0092), YIB (*F*_ST_ = 0.0100), ENS (*F*_ST_ = 0.0128), FUL (*F*_ST_ = 0.0129), YAN (*F*_ST_ = 0.0148), DEC (*F*_ST_ = 0.0189), XIL (*F*_ST_ = 0.0229), and DEH (*F*_ST_ = 0.0498). For the river group, crossbreds showed a close relationship with the Murrah (*F*_ST_ = 0.0542) and Nili-Ravi (*F*_ST_ = 0.0724) breeds.

### Linkage disequilibrium and Autozygosity Segments

Overall estimated LD between river and swamp groups was different in the present study (**Fig 3A**). Compared to the river group (*r*^*2*^ =0.61), the highest maximum average LD (*r*^*2*^=0.88) was found within the swamp group. As expected, LD declined as the physical distance between pairwise SNPs. LD decay in swamp group had declined markedly compared with the river group. The average distance when LD decay dropped to the value of 0.2 was approximately 15 Kb for swamp and 50 Kb for river group, respectively.

**Fig 3.**
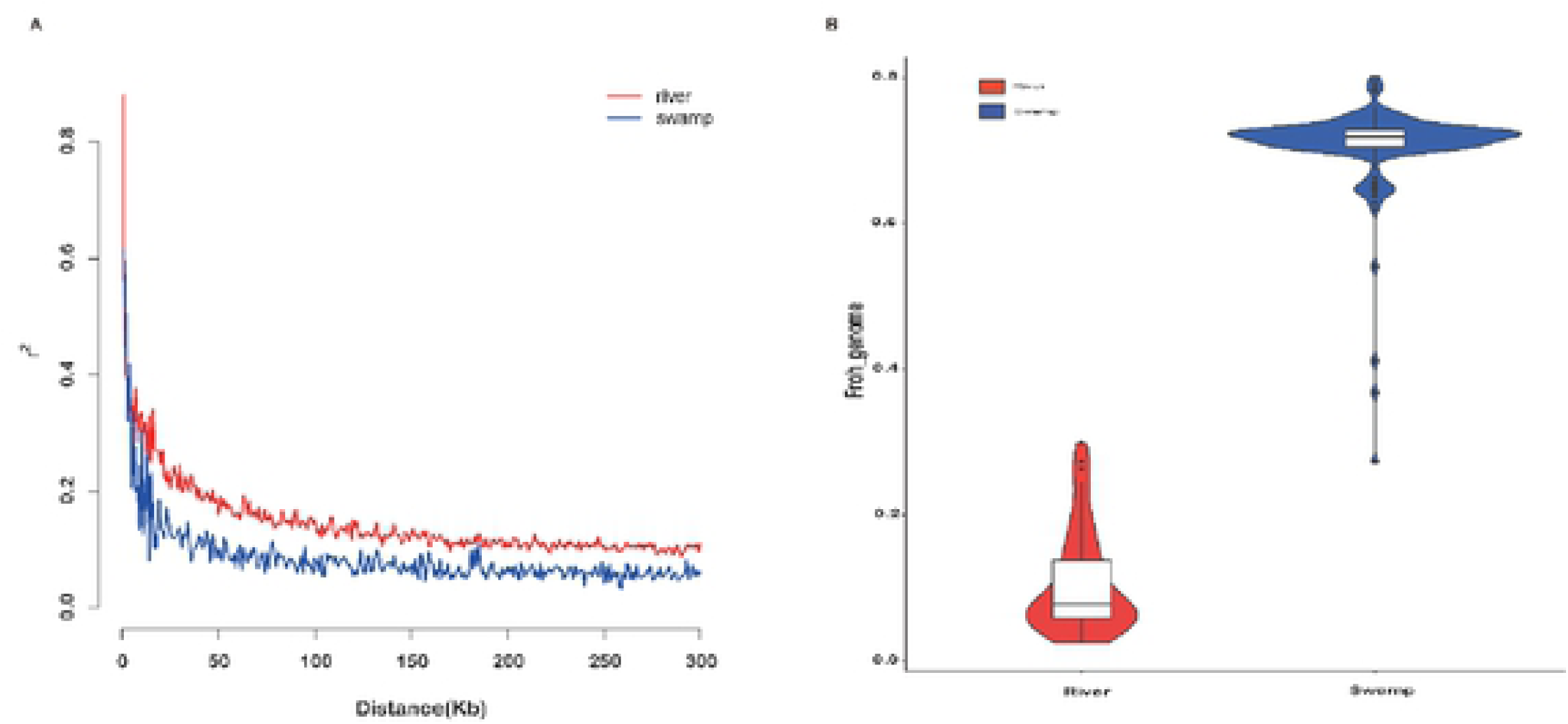
LD decay and autozygosity frequency distribution of ROH between river and swamp groups.

To estimate the recent inbreeding, we performed the genome-wide autozygosity analysis with the runs of homozygosity (ROH) between river and swamp groups. The result showed that buffaloes in swamp group had higher overall levels of F_ROH_ than that of river group (**Fig 3B**). Moreover, we also estimated the ROH distribution by the length between the river and swamp groups (**S1 Fig**). The results showed that the difference in genetic diversity was found between river and swamp group. Buffaloes in swamp group had the lower fraction of ROH in short tract (0-2 Mb), while the river breed exhibit a higher faction of ROH in long tract (>16 Mb).

### Signatures of selection

The hapFLK statistic that accounted for the haplotype information and hierarchical structure [36, 37] was used to identify selection footprints on the contrast model (river *vs.* swamp). HapFLK analyses revealed that a total of 12 genomic regions was identifed (**Fig 4**). These regions were located on chromosomes 1 (197,939,132-201,746,066 Kb), 2 (84,613,461-91,314,070 Kb), 8 (80,317,117-88,124,162 and 96,080,274-98,588,633 Kb), 11 (25,584,886-27,617,098 Kb), 12 (15,979,826-16,215,148 Kb), 15 (7,435,513-13,248,436 Kb), 16 (64,367,119-69,995,014, 70,024,989-79,994,535, and 80,043,731-83,241,865 Kb), 19 (69,537,494-71,631,407 Kb), and 24 (12,608,124-13,746,779 Kb). **Table 2** summarizes the annotation information of outlier SNPs within the contrast model. The candidate selective sweep regions ranged from 0.24 Mb to 9.97 Mb on Chr12 and Chr16, respectively.

**Table 2.** Summary of the selective sweep regions detected using hapFLK analyses between river and swamp groups. CHR, chromosome.

| CHR | Start (kb) | End (kb) | Genomic region (Mb) | Number of candidate genes | Top significant SNP | Genes mapping top SNPs |
| --- | --- | --- | --- | --- | --- | --- |
| 1 | 197939132 | 201746066 | 3.81 | 15 | AX-85111638 | KCNH8 |
| 2 | 84613461 | 91314070 | 6.70 | 41 | AX-85092726 | ACVR1C |
| 8 | 80317117 | 88124162 | 7.81 | 31 | AX-85062676 | HYAL4,SPAM1 |
| 8 | 96080274 | 98588633 | 2.51 | 14 | AX-85063572 | CHCHD3 |
| 11 | 25584886 | 27617098 | 2.03 | 32 | AX-85056732 | - |
| 12 | 15979826 | 16215148 | 0.24 | 3 | AX-85129983 | - |
| 15 | 7435513 | 13248436 | 5.81 | 35 | AX-85085538 | DEPTOR |
| 16 | 64367119 | 69995014 | 5.63 | 27 | AX-85069732 | LOC112579725 |
| 16 | 70024989 | 79994535 | 9.97 | 25 | AX-85130034 | LOC112579725 |
| 16 | 80043731 | 83241865 | 3.20 | 9 | AX-85085883 | - |
| 19 | 69537494 | 71631407 | 2.09 | 31 | AX-85108518 | TPPP |
| 24 | 12608124 | 13746779 | 1.14 | 3 | AX-85085586 | CALN1 |
| Total |  |  | 4.24 |  |  |  |

**Fig 4.**
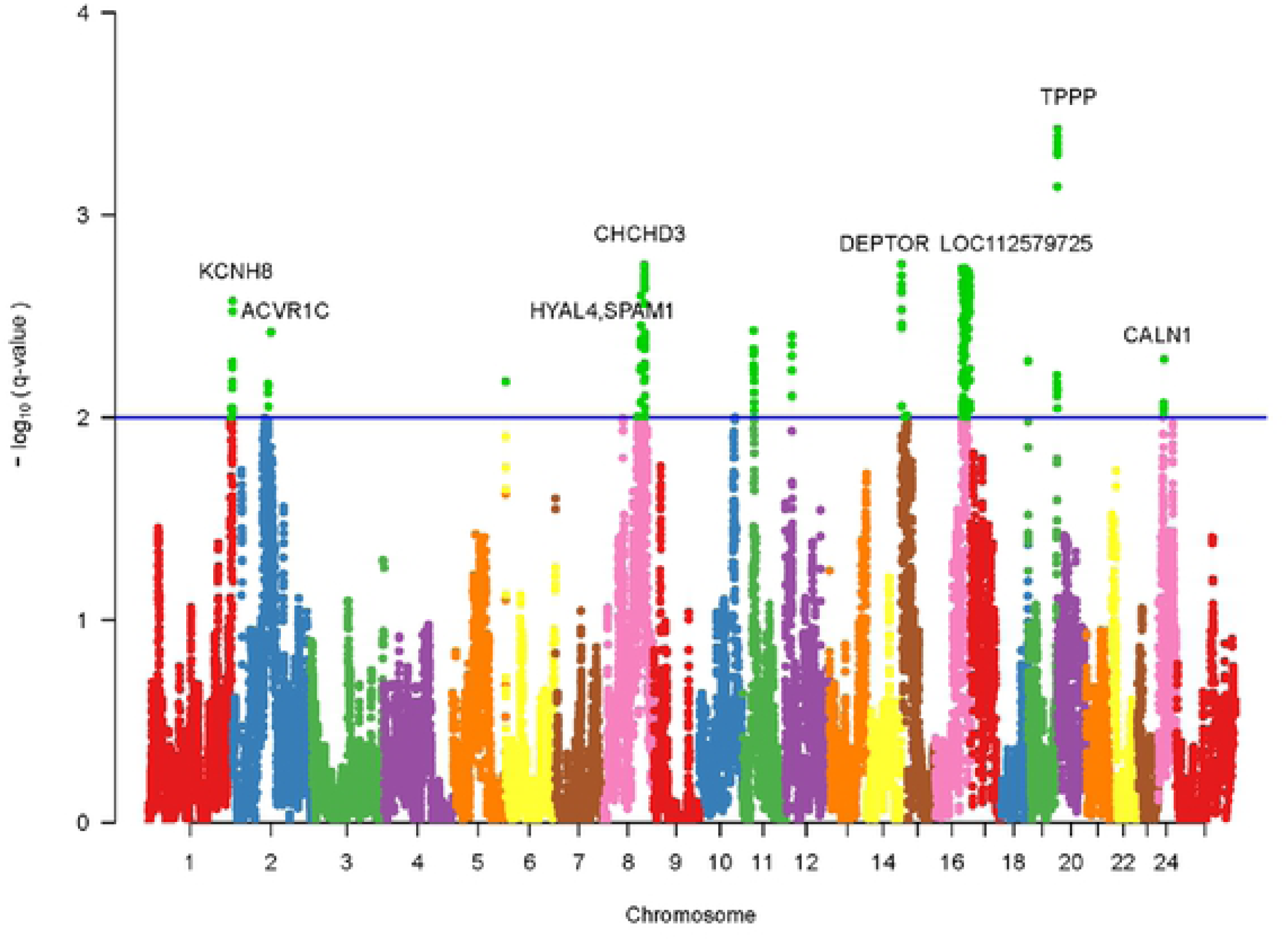
Whole-genome scan for selective sweeps using hapFLK statistic. The blue line represents a significant threshold.

The average length of the candidate regions was 4.24 Mb. The largest number of SNPs (35) within a genomic region was found in Chr15 with a length of 5.81 Mb. Notably, a total of 12 top significantly SNPs was identified, corresponding to the 8 candidate genes and one lncRNA (LOC112579725) fragment.

### Identification of QTLs

To further identify the QTLs associated with milk production traits in buffaloes, we firstly performed the haplotypes analysis using a 0.5-Mb window around the top significantly SNPs (**Fig 5**). The results showed that a total of 18 blocks was identified and located on chromosomes 8, 11, 12, 16, 19, and 24, respectively. For them, a total of 4 blocks (11_Block2, 12_Block2, 16_Block1, and 19_Block5) was found to have the top significantly SNPs. The largest length of blocks was the 19_Block5 with the length of 264 Kb. The haplotype association analysis revealed that a total of 12 blocks was identified to be associated with milk production traits (**Table 3**). Interestingly, 3 blocks (11_Block2, 19_Block3, and 19_Block4) had significantly genetic effects on all milk traits in buffaloes (*P*<0.05). The 8_Block2, 12_Block2, and 19_Block1 were shown to be associated with milk protein percentage (PP) and fat percentage (FP) in buffaloes (*P*<0.05). Moreover, we found that a total of 6 blocks (8_Block3, 12_Block1, 16_Block1, 19_Block2, 19_Block5, and 24_Block1) were associated with milk yield (MY), fat yield (FY) or protein yield (PY) in buffaloes (*P*<0.05).

**Table 3.**
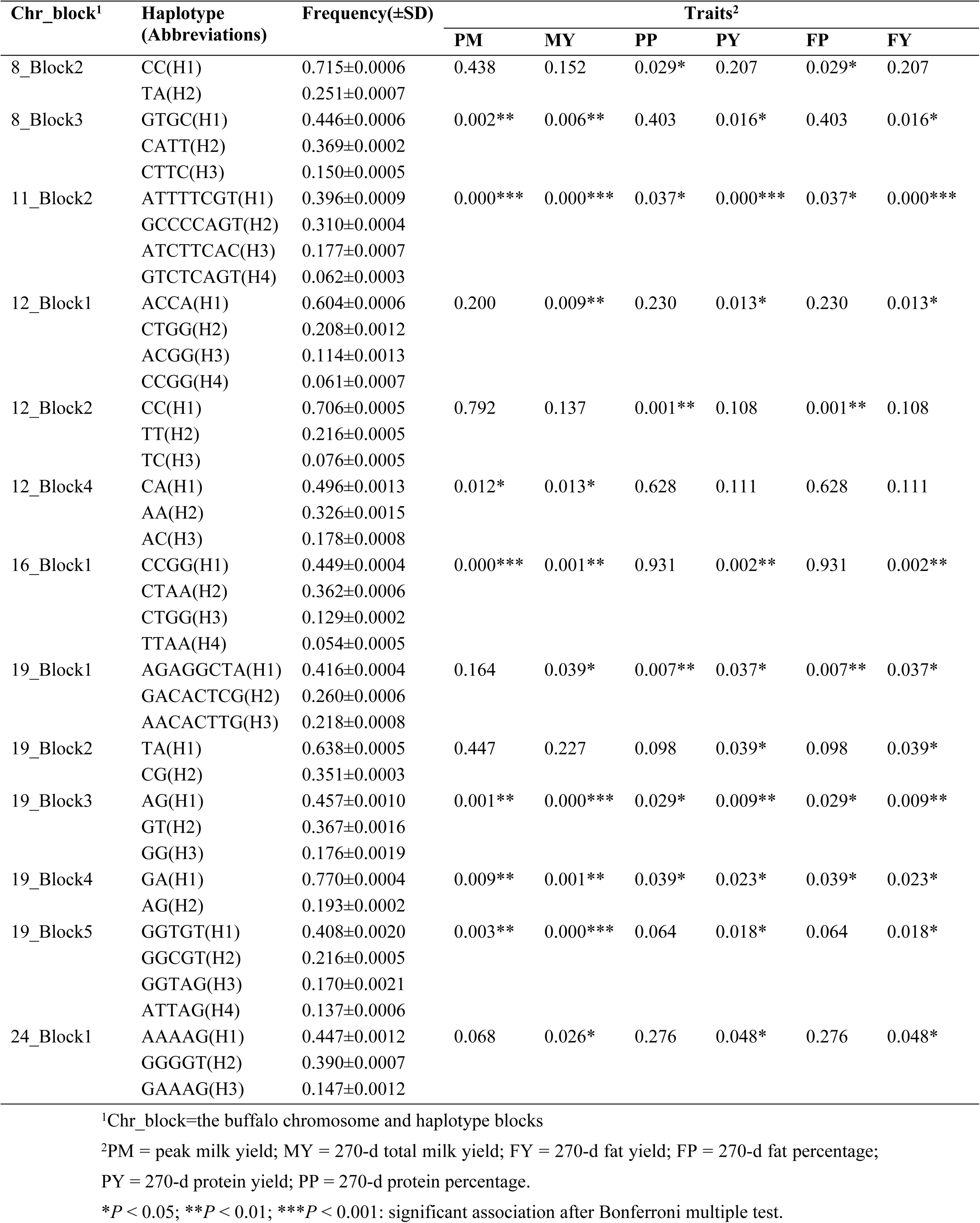
Haplotype association analyses for the milk production traits in buffaloes.

**Fig 5.**
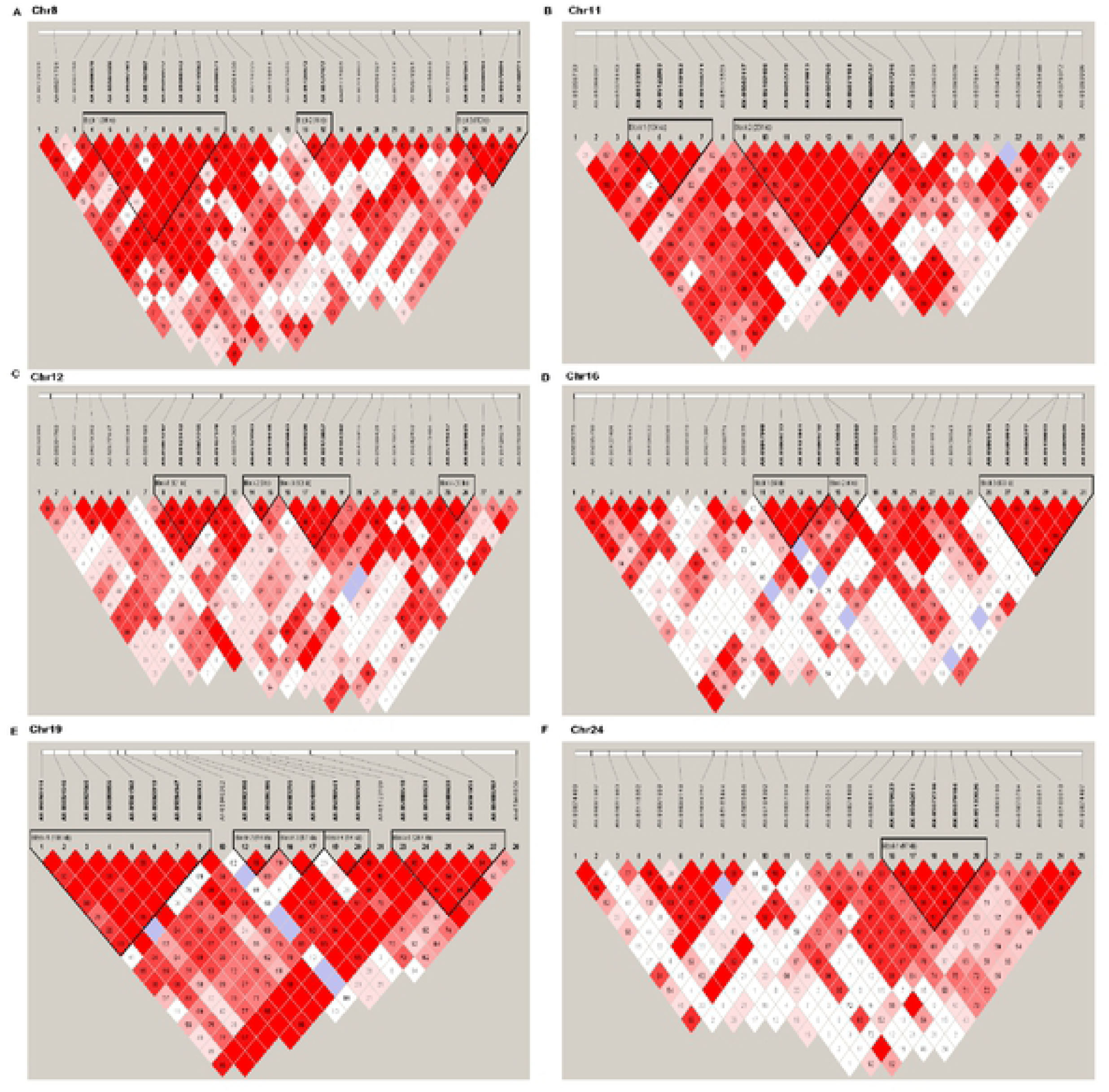
Haplotype patterns for the significant SNP based on LD within detected regions.

For the blocks harbored the top significantly SNPs, we further performed the haplotype combination analysis in the present study (**Table 4)**. Bonferroni analysis revealed that buffaloes with diplotype H2H2 in 12_Block2 were highest significantly associated with FP and PP than the other diplotype (*P* < 0.05). In the 11_Block2, the individuals with diplotype H1H3 showed a higher MY, FY, and PY compared to other diplotypes, while the diplotype H3H4 and H1H2 had the highest peak milk yield (PM) and fat or protein percentages (*P* < 0.05), respectively. For 16_Block1, buffaloes with diplotype H2H2 had highest significantly associated with PM, FY, and PY, while the diplotype H1H4 exhibited the highest genetic effect on MY (*P* < 0.05). Moreover, these buffaloes with the diplotype H2H4 in 19_Block5 displayed a higher PM, MY, FY, and PY compared to other diplotypes (*P* < 0.05).

**Table 4.** Genetic effects of haplotype combinations among the four QTL regions on milk production traits in buffaloes.

| Blocks | Haplotype combination | Traits |  |  |  |  |  |
| --- | --- | --- | --- | --- | --- | --- | --- |
| | | PM(±SD) | MY(±SD) | FP(±SD) | PP(±SD) | FY(±SD) | PY(±SD) |
| 11_Block2 | H1H1 | 15.53±0.43ab | 2918.31±87.94b | 7.96±0.14a | 7.96±0.14a | 231.32±7.75b | 231.32±7.75b |
| | H1H2 | 15.28±0.40ab | 2850.28±84.76bc | 8.03±0.14a | 8.03±0.14a | 227.12±7.48b | 227.12±7.48b |
| | H1H3 | 16.45±0.44a | 3147.73±92.30a | 7.94±0.15a | 7.94±0.15a | 248.41±8.14a | 248.41±8.14a |
| | H1H4 | 16.28±0.43a | 3038.39±91.12a | 8.03±0.15a | 8.03±0.15a | 243.68±8.03a | 243.68±8.03a |
| | H2H2 | 15.70±0.44a | 2975.41±92.62a | 7.72±0.15b | 7.72±0.15b | 228.92±8.17bc | 228.92±8.17bc |
| | H3H2 | 16.31±0.42a | 3053.66±88.38a | 7.90±0.14a | 7.90±0.14a | 241.38±7.78a | 241.38±7.78a |
| | H3H3 | 15.24±0.54ab | 3008.24±114.43a | 8.09±0.19a | 8.09±0.19a | 242.34±10.08a | 242.34±10.08a |
| | H3H4 | 16.99±0.65a | 3174.30±136.78a | 7.98±0.22a | 7.98±0.22a | 253.45±12.04a | 253.45±12.04a |
| | H4H2 | 15.87±0.59a | 3025.39±124.84a | 8.14±0.20a | 8.14±0.20a | 244.27±10.99a | 244.27±10.99a |
| | H4H4 | 15.79±0.86a | 2914.81±182.73a | 8.03±0.30a | 8.03±0.30a | 232.10±16.09 | 232.10±16.09a |
| 12_Block2 | H1H1 | 15.75±0.39 | 2992.38±83.00 | 7.91±0.14ab | 7.91±0.14ab | 235.53±7.23 | 235.53±7.23 |
| | H2H1 | 15.76±0.39 | 2943.13±83.01 | 8.04±0.14a | 8.04±0.14a | 236.14±7.23 | 236.14±7.23 |
| | H2H2 | 15.83±0.50 | 2858.51±105.84 | 8.31±0.17a | 8.31±0.17a | 237.00±9.21 | 237.00±9.21 |
| | H2H3 | 15.30±0.51 | 2830.18±107.76 | 7.96±0.18a | 7.96±0.18a | 222.73±9.38 | 222.73±9.38 |
| | H3H1 | 15.76±0.43 | 2919.22±91.28 | 7.79±0.15ab | 7.79±0.15ab | 226.92±7.95 | 226.92±7.95 |
| | H3H3 | 14.67±1.04 | 2844.60±218.48 | 7.53±0.36a | 7.53±0.36a | 211.77±19.01 | 211.77±19.01 |
| 16_Block1 | H1H1 | 15.55±0.39ab | 2944.88±83.37a | 7.97±0.14 | 7.97±0.14 | 233.73±7.28a | 233.73±7.28a |
| | H1H2 | 15.50±0.41ab | 2889.12±86.01ab | 7.97±0.14 | 7.97±0.14 | 229.57±7.50ab | 229.57±7.50ab |
| | H1H3 | 15.34±0.42ab | 2846.89±88.02ab | 7.94±0.15 | 7.94±0.15 | 225.29±7.67ab | 225.29±7.67ab |
| | H1H4 | 16.34±0.46a | 3083.19±96.24a | 7.94±0.16 | 7.94±0.14 | 242.76±8.39a | 242.79±8.39a |
| | H2H2 | 16.36±0.42a | 3044.85±88.00a | 7.96±0.15 | 7.96±0.15 | 242.96±7.67a | 242.96±7.67a |
| | H2H4 | 16.01±0.52a | 2840.34±110.07a | 8.08±0.18 | 8.08±0.18 | 229.46±9.65a | 229.46±9.65a |
| | H3H2 | 15.59±0.46a | 2970.29±96.50a | 7.99±0.16 | 7.99±0.16 | 237.52±8.41a | 237.52±8.41a |
| | H3H3 | 15.30±0.64a | 2842.73±136.18a | 7.81±0.23 | 7.81±0.23 | 219.17±11.87a | 219.17±11.86a |
| | H3H4 | 14.56±0.91a | 2754.95±192.82a | 7.68±0.32 | 7.68±0.32 | 212.21±16.80a | 212.21±16.80a |
| 19_Block5 | H1H1 | 15.57±0.41ab | 2917.36±85.48b | 8.04±0.14 | 8.04±0.14 | 233.27±7.57a | 233.26±7.57a |
| | H1H2 | 15.84±0.42a | 2991.74±86.46a | 7.96±0.15 | 7.96±0.15 | 237.28±7.64a | 237.28±7.64a |
| | H1H3 | 15.68±0.40ab | 2907.34±83.41b | 7.96±0.14 | 7.96±0.14 | 230.68±7.37a | 230.68±7.38a |
| | H1H4 | 15.47±0.43ab | 2969.81±89.94b | 7.99±0.15 | 7.99±0.15 | 235.78±7.95a | 235.78±7.95a |
| | H2H2 | 15.81±0.48a | 2967.22±99.91b | 8.12±0.17 | 8.12±0.17 | 240.37±8.83a | 240.37±8.83a |
| | H2H4 | 16.55±0.45a | 3195.95±93.87a | 7.72±0.16 | 7.72±0.16 | 246.58±8.30a | 246.58±8.30a |
| | H3H2 | 15.68±0.44a | 2962.21±91.49b | 8.01±0.15 | 8.01±0.15 | 235.94±8.08a | 235.94±8.08a |
| | H3H3 | 15.87±0.62a | 2996.27±128.29a | 7.95±0.22 | 7.95±0.22 | 237.81±11.49a | 237.81±11.49a |
| | H3H4 | 15.90±0.53a | 2893.09±109.42b | 7.70±0.18 | 7.70±0.18 | 223.73±9.66a | 223.73±9.66a |
| | H4H4 | 13.80±0.68ab | 2586.08±141.34b | 8.10±0.24 | 8.10±0.24 | 205.68±12.75ab | 205.68±12.75ab |

As shown in **Table 5**, a total of 7 genes and 1 lncRNA were predicted. For them, the *SLC9A3, EXOC3, AHRR, CEP72, PRB1*, and *TPPP* genes in 19_Block5, while *FUT8* and LOC112587805 in 11_Block2, were served as the candidate genes associated with buffalo milk production traits. However, no candidate gene was discovered within the 12_Block2 and 16_Block1 regions.

**Table 5.**
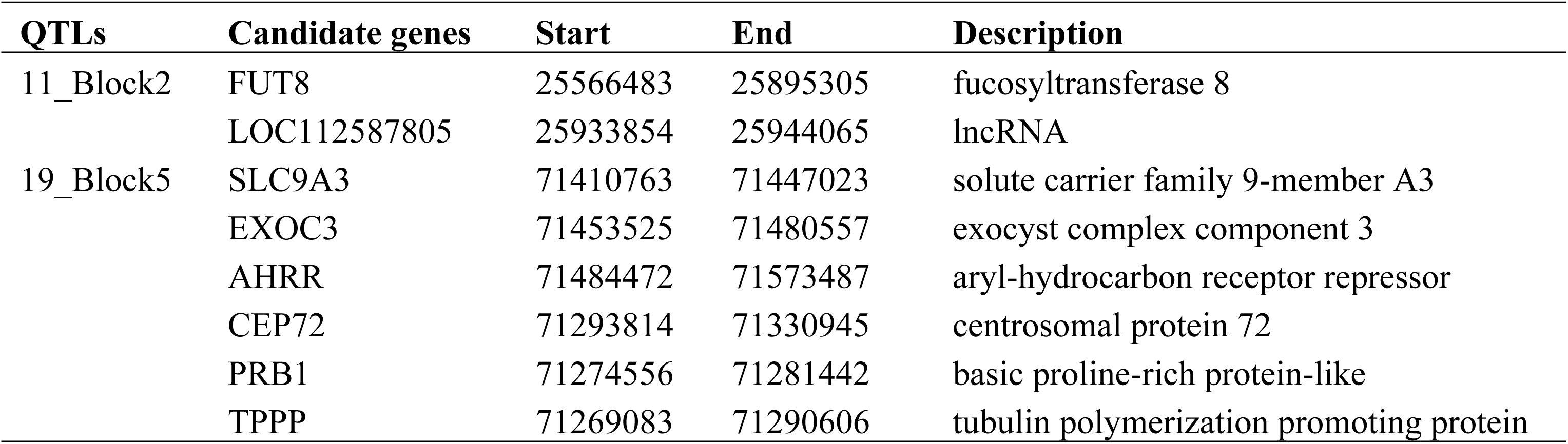
Details of candidate genes within the two QTL regions.

| QTLs | Candidate genes | Start | End | Description |
| --- | --- | --- | --- | --- |
| 11_Block2 | FUT8 | 25566483 | 25895305 | fucosyltransferase 8 |
|  | LOC112587805 | 25933854 | 25944065 | lncRNA |
| 19_Block5 | SLC9A3 | 71410763 | 71447023 | solute carrier family 9-member A3 |
|  | EXOC3 | 71453525 | 71480557 | exocyst complex component 3 |
|  | AHRR | 71484472 | 71573487 | aryl-hydrocarbon receptor repressor |
|  | CEP72 | 71293814 | 71330945 | centrosomal protein 72 |
|  | PRB1 | 71274556 | 71281442 | basic proline-rich protein-like |
|  | TPPP | 71269083 | 71290606 | tubulin polymerization promoting protein |

## Discussion

China is rich in buffalo resources that are mainly distributed in 18 provinces in the central and south regions of the country. The Chinese buffalo breeds have a lower milk production in comparison with river buffalo breeds. In this work, we performed a population analysis of thirteen buffalo breeds in South China. PCA and phylogenetic analysis both showed that two distinct clusters formed without overlap among the studied breeds; ten indigenous Chinese buffaloes were clustered into the swamp group, Murrah and Nili-Ravi breeds were grouped into the river group, while the crossbred was concordantly located between the two groups. Population structures analysis further showed that there was significant admixture, showing that the breeds are genetically distinct between river and swamp groups. In this regard, we further investigated the genetic diversity, LD and ROH, signatures of selection, and identification of QTLs on the contrast model: the river (high milk production) *vs.* swamp (low milk production) groups. These help in the understanding of the genetic basis of milk production traits in dairy buffaloes, which promote the development of dairy buffalo industry.

Using the medium density SNP chip data, we found that buffalo breeds in swamp group had lower genetic diversity than that of the river group based on heterozygosity measures. Similar results were also reported by Colli et al. [10]. In fact, Chinese indigenous breeds sustain higher levels of genetic variability than river breeds. The assumption was supported by previous reports that utilized microsatellite markers [38–40] and mtDNA [41, 42]. This can be explained by the current buffalo 90K array is optimized for use in four river buffalo breeds (Mediterranean, Murrah, Nili-Ravi, and Jaffarabadi buffaloes) and has the lower representation of swamp breeds [43]. The disproportionate distribution of MAF between river and swamp groups is another reason to influence the SNP ascertainment bias. For example, buffalo breeds in swamp group had the highest proportion of SNPs with lower MAF category compared to the river group. The relatively high proportion of SNPs with high MAF in the Chinese crossbred buffalo can be attributed to the fact that these buffaloes were recent crossbreeds with a significantly high inheritance of Murrah and Nili-Ravi breeds ancestry. Moreover, we also observed differences in the genetic differentiation among the studied breeds. As expected *F*_ST_ among the swamp breeds was lower than the river and crossbred breeds, ranging from 0.0076 to 0.7535, suggesting a lack of differentiation among Chinese swamp breeds. This was lower than 0.52 observed among the African buffalo (*Syncerus caffer*) breeds by Smitz, Berthouly (44), but higher than the *F*_ST_ estimates reported in previous studies [45–47]. These differences may be caused by the following: 1) small population effect sizes that may be resulted in data error, 2) differences in buffalo breeds or geographical distribution, and 3) differences in the estimated methods or markers density.

Analysis of the genome-wide LD decay plays a vital role in the GWAS mapping of loci associated with economically important traits in livestock animals. Our previous studies have demonstrated that the LD extent was different between purebred and crossbred buffalo populations, with purebred having highest levels of LD [5]. In this study, we estimated the LD extent between river and swamp groups, showing that the LD decay in swamp group had declined markedly compared with the river group. This result suggested that these buffaloes in swamp group had a higher genetic diversity. Of note, LD extent across populations is much shorter in Chinese swamp buffaloes than the river and crossbred buffaloes. The average distance over which LD decay dropped to the value of 0.2 was approximately 15 Kb for swamp and 50 Kb for river group, respectively. This finding was also supported by previous studies [5]. Moreover, it is well known that the Chinese indigenous breeds underwent a longer time of domestication compared with the river breeds. The assumption was also supported by the results from ROH analysis. The buffaloes in swamp group had the lower fraction of ROH in short tract (0-2 Mb), while the river breed exhibit a higher faction of ROH in long tract (>16 Mb). Interestingly, evidence indicated that ROH could be used to assist with the interpretation of the inbreeding coefficient and give insights about populations history [48, 49]; the short and long ROH can respectively reflect the remote and recent inbreeding activity [50]. Apparently, our data showed that the extensive remote and recent inbreeding activity was found in the swamp and river group, respectively.

Understanding the signatures of selection detected within livestock breeds can contribute to the identification of genomic regions that are or have been targeted by selection [51]. Here, we performed the selective sweeps analysis between river and swamp groups using hapFLK analysis. A total of 12 genomic regions was identified with an average length of 4.24 Mb. In these regions, we found that a total of 12 top significantly SNPs detected within 9 chromosomes, corresponding to 8 candidate genes and 1 lncRNA (LOC112579725) fragment. Most of the candidate genes were predicted to be associated with cell structure and function. For example, *CHCHD3*, as the inner mitochondrial membrane scaffold protein, plays a role in cell growth and oxygen consumption [52]. *HYAL4* and its paralog gene *SPAM1* has demonstrated to be similar in structure to hyaluronidases that were one of the major glycosaminoglycans of the extracellular matrix involved in cell proliferation, migration, and differentiation [53–55]. *ACVR1C* is a type I receptor for the TGFB family of signaling molecules that plays a role in cell differentiation, growth arrest, and apoptosis [56]. Numerous studies have demonstrated that *DEPTOR* is the negative regulator of the mTORC1 and mTORC2 signaling pathways, which play a vital role in cell growth, metabolism, and disease [57, 58]. *TPPP* has confirmed to play a role in the polymerization of tubulin into microtubules that involved in the multiple biological functions such as the cell migration, cilia and flagella, development, and gene regulation [58–60].

QTL mapping is critical for the gene cloning, molecular-marker-assisted selection breeding, and trait improvement, which has been widely used for animal [61] and plant breeding [62]. To further identify the QTLs associated with milk production traits in buffaloes, we performed the haplotypes analysis using a 0.5-Mb window around the top significantly SNPs. To our acknowledgment, information on the identification of QTLs associated with milk production traits in buffalo is limited. Liu et al. [63] found that 2 genomic regions were found to associate with buffalo milk production traits. Here, we discovered a total of 18 blocks within 6 chromosomes. Among these blocks, 13 blocks were found to be significantly associated with the milk production traits in buffaloes. Interestingly, 4 blocks (11_Block2, 12_Block2, 16_Block1, and 19_Block5) harbored the top significantly SNPs have also been significantly associated with milk production traits in buffaloes (*P* <0.05). Notably, the diplotype H2H2 in 12_Block2, H1H3 in 11_Block2, H2H2 in 16_Block1, and H2H4 in 19_Block5 can consider as the dominant haplotype combinations related to milk traits in buffaloes based on the Bonferroni analysis. For these QTL regions, moreover, we found that some candidate genes can be predicted in the 11_Block2 and 19_Block5, while no candidate gene was discovered within the 12_Block2 and 16_Block1 regions. For the 11_Block2, the *FUT8* was found to be a signaling receptor involved in many physiological and pathological processes [64], implying that this QTL might be related to cell growth. In the 19_Block5, the *SLC9A3, EXOC3, AHRR, CEP72, PRB1*, and *TPPP* genes were predicted. As the *AHRR* participated in the aryl hydrocarbon receptor (AhR) signaling cascade [65], *CEP72* [66] and *TPPP* [67] participated in the microtubule formation; the result suggested that these genes might be related to the cell growth and differentiation. Notably, Basham et al. [68] reportd that *AHRR* could mediate the AHR singnal patwahy to block milk production in mammary epithelial cell and block transcription of the milk gene *β*-casein. *SLC9A3* was the member of solute carrier family that played a vital role in the signal transduction, and amino acid as well as glucose transporter [69]. Although limited information on these genes associated with milk production traits has been reported, we have reason to assume that these candidate genes should have genetic effects on milk production traits based on their biological function. Further research of these candidate genes is warranted to explore the underlying molecular mechanism of the phenotypic traits in buffaloes.

## Materials and methods

### Ethics Statement

All animal work, experimental protocols, and animal care were approved by the Animal Ethics Committee of the Buffalo Research Institute (BRI), Chinese Academy of Agricultural Sciences (CAAS) (approval code GXBRI-06-2019).

### Sampling and Genotyping

A total of 176 unrelated buffaloes representing 13 breeds were included in the present study (**Table 1**), and their geographical origin was shown in **Fig 6**. We collected a total of 102 blood samples, of which 23 river and 20 crossbred buffaloes were from BRI-CAAS, and 59 swamp buffaloes were collected from different villages in Southwest China. Moreover, we obtained publicly available genotypic data from 74 buffaloes provided by Colli et al. [10].

**Fig 6.**
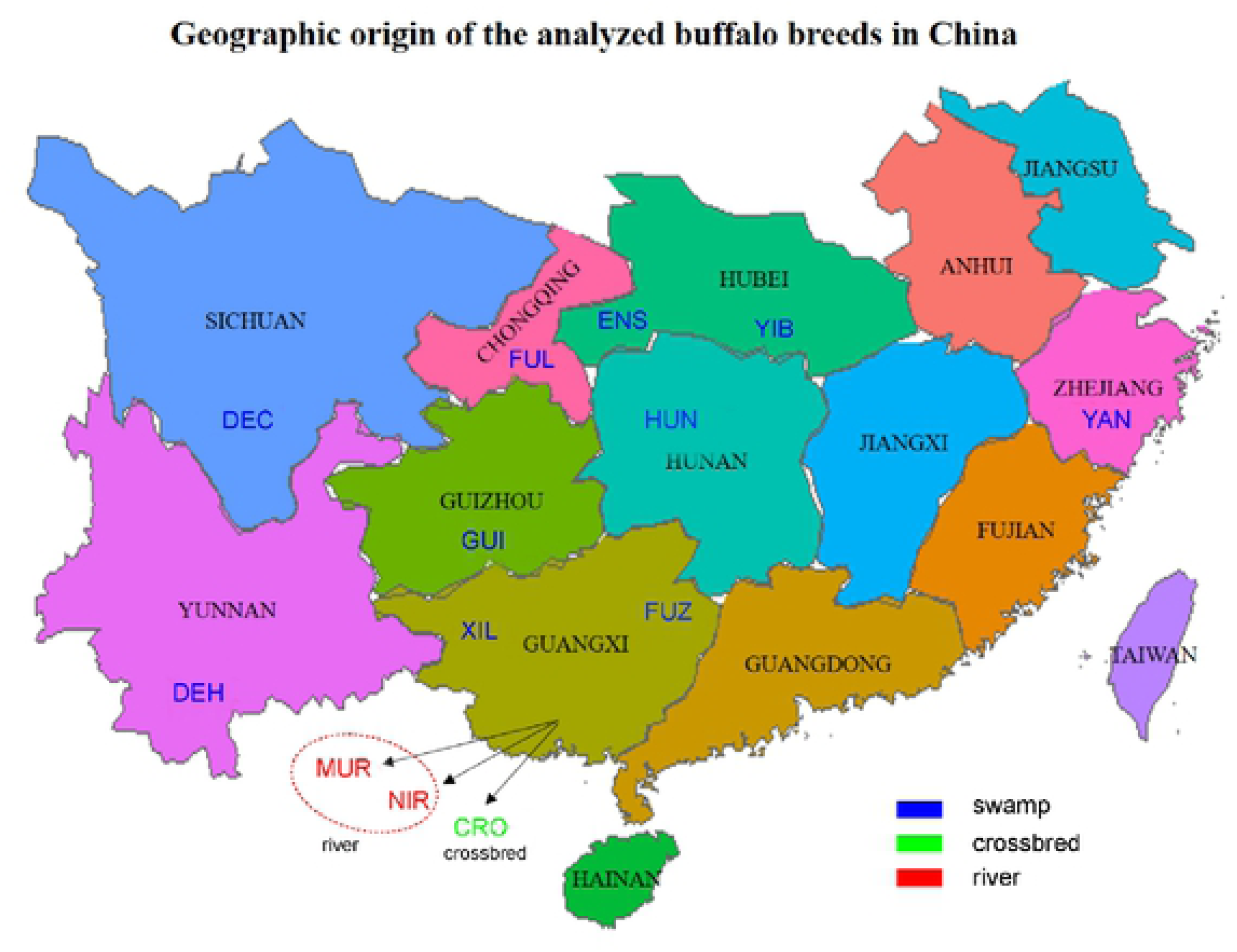
Geographic origin of the analyzed buffalo breeds in South China region. Note: The R software [80] with the maptools and ggplot2 packages were used to create the map and corresponded to the database was downloaded from the National Geomatics Center of China (http://www.ngcc.cn/).

Genomic DNA for each blood sample was isolated using the TIANamp Blood DNA Kit [Tiangen Biotech (Beijing) Co., Ltd., Beijing, China]. Quality and quantity of isolated DNA were detected using the NanoDrop2000 spectrophotometer (Thermo Fisher Scientific, Wilmington, DE, USA) and 1.5% Gel electrophoresis, respectively. SNP genotyping was performed using the Axiom^®^ Buffalo Genotyping Array (Affymetrix, Santa Clara, CA, USA). For the quality control, SNPs with MAF ≤ 0.05, SNP call rate ≤ 95%, individual call rate ≤ 95% were excluded using the PLINK 1.9 [70]. After filtering the duplicate and unknown chromosomal SNPs, a total of 35,547 SNPs was used for subsequent analysis.

### Population genetics analysis

To explored the individuals’ relationship in the analyzed breeds, we determined the eigenvectors significance using EIGENSOFT 7.2 [71] software with the Tracey-Widom test and visualized the PCA plot with in-house R scripts. PHYLIP 3.697 [72] was conducted to calculate the distance matrix, and MEGA7 [73] was used to present the neighbor-joining tree. Population structure among the analyzed buffaloes was determined using the ADMIXTURE 1.3.0 [74] software with the cluster (K) number set from 2 to 4. Admixture plots were further visualized using the Microsoft Office Excel 2016.

### Genetic diversity

Genome-wide genetic variability within buffalo breeds was measured by the Arlequin 3.5.2 [75] software with the three metrics: H_O_, He, and A_R_. Population differentiation for the pairwise genetic differentiation *F*_*ST*_ value was measured using the adegenet [76] R-package.

### Linkage disequilibrium decay

Genome-wide LD was evaluated between river and swamp groups. To diminish the SNP ascertainment bias, we randomly selected the same sample numbers (n=43) for each group. For all pairs of SNPs, the pair-wise LD between river and swamp groups was calculated and visualized using the PopLDdecay [77].

### Identification of Runs of Homozygosity

ROH between river and swamp groups was detected using the sliding-window based method implemented in detectRUNS [78] R-package with the following parameters: windowSize=15, threshold=0.05, minSNP=20, and minDensity=1/50000. F_ROH_ on the contrast model was calculated using the following equation:

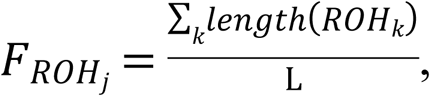

where *ROH*_*k*_ = the *k*th ROH in individual *j*’s genome and L = the total length of the genome (or X-chromosome).

### Identification of selection signatures

Selection signature analysis on the contrast model was performed using the hapFLK 1.4 [79] software. Briefly, we firstly performed LD pruning analysis and admixture analysis to obtain the population IDs for hapFLK analysis, which was respectively implemented in the PLINK1.9 and ADMIXTURE 1.3.0. The main parameters for LD pruning analysis were as follows: a window size of 50 SNPs, a step size of 5 bp and an *r*^*2*^ threshold of 0.5. In the hapFLK analysis, the number of clusters (-K) was set to 2, and the expectation maximization iterations (--nfit) were specified to 20. The hapFLK values were adjusted using the following equation:

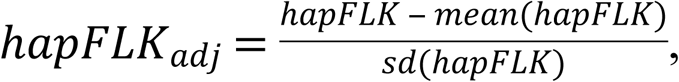

where the mean and sd values of hapFLK were calculated using the MASS R-package. The q-values of hapFLK_adj_ was computed using a chi-square distribution with in-house R scripts. A q-value threshold of 0.01 was applied to limit the number of false positives.

### Haplotype and association analysis

Phenotypic and genotypic data for 489 Italian Mediterranean buffaloes was provided by Liu et al. [63]. Phenotypic data for the peak milk yield (PM), total milk yield (MY), fat yield (FY), fat percentage (FP), protein yield (PY), and protein percentage (PP) was included during a 13-yr period (2002-2014). The identified genomic regions were surveyed to construct haplotype blocks within 0.5 Mb of the top significantly SNPs using HaploView 4.2 (Barrett et al., 2005). Association between each haplotype combination and 6 milk production traits were performed using the SAS 9.4 (SAS Institute Inc., Cary, NC) software with the following model:

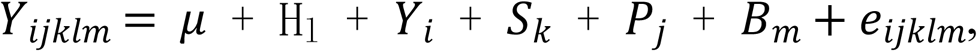

where *Y*_*ijklm*_ = the trait observation, *μ* = the overall mean, H_l_ = the fixed effect of the *l*th herd (4 farms), *Y*_*i*_ = the fixed effect of the *i*th year, *S*_*k*_ = the fixed effect of the *k*th season of calving (2 seasons), *P*_*j*_ = the fixed effect of the *j*th parity classes (1 to 7 and ≥8), *B*_*m*_ = the fixed effect of the *m*th haplotype block, and *e*_*ijklm*_ = the random residual. Bonferroni t-test for pairwise comparisons among different levels of fixed effects was used. The threshold of *P*-value < 0.05 was used to identify the haplotype blocks affecting buffalo milk traits identified as the candidate QTLs.

### Gene Annotation

We annotated the identified regions under significant selection pressure using the NCBI’s Genome Data Viewer on the buffalo genome (*UOA_WB_1*). Genes within a region spanning 50 Kb upstream and downstream of the candidate selection regions were annotated. Moreover, the identification of candidate genes within the QTLs was also performed based on the current buffalo genome.

## Supporting information captions

**S1 Fig. Distribution of autosomal ROH between river and swamp groups**

**S1 Table. Distribution of SNP number per chromosome, mean length (Mb), mean length of adjacent SNPs and average LD for the markers set after quality control**

**S2 Table. Estimates of the pairwise genetic differentiation statistic among breeds (*F***_***ST***_ **statistics; below the diagonal).**

**S1 Data Sheets. Genotype datasets of 176 buffaloes were used in this study.**

## Acknowledgments

We thank the anonymous for the smallholders for their supports on the buffalo samples.

## Funding Statement

Funded by the Major Science and Technology Projects in Guangxi (AA16450002), Central Guidance for Local Science and Technology Development Projects (ZY18164003), and Natural Science Foundation of Guangxi (2017GXNSFBA198191 and 2017GXNSFBA198022). The funders had no role in study design, data collection and analysis, decision to publish, or preparation of the manuscript.

## Author Contributions

**Conceptualization:** Ting-Xian Deng

**Formal analysis:** Ting-Xian Deng

**Funding Acquisition:** Ting-Xian Deng and Xian-Wei Liang

**Investigation:** Xing-Rong Lu, An-Qin Duan, Xiao-Ya Ma, Sha-Sha Liang

**Methodology:** Ting-Xian Deng, Xing-Rong Lu, and An-Qin Duan

**Resources:** Ting-Xian Deng and Xian-Wei Liang

**Writing-original draft:** Ting-Xian Deng

**Writing-review & editing:** Ting-Xian Deng and Xing-Rong Lu

## Data Availability

The genotype datasets of this study are provided as the Supplementary Data Sheets S1.

## References

1. Mishra B, Dubey P, Prakash B, Kathiravan P, Goyal S, Sadana D, et al. Genetic analysis of river, swamp and hybrid buffaloes of north-east India throw new light on phylogeography of water buffalo (Bubalus bubalis). Journal of Animal Breeding and Genetics. 2015;132(6):454–66.

2. Cui B, Wang F, Cui Y, Zhang Y, Li J, Hui T, et al. A bried analysis of the current status of the Chinese buffalo industry. Meat research. 2013;27:37–40.

3. Degrandi TM, Pita S, Panzera Y, Oliveira EHCD, Marques JRF, Figueiró MR, et al. Karyotypic evolution of ribosomal sites in buffalo subspecies and their crossbreed. Genetics & Molecular Biology. 2014;37(2):375–80.

4. Harisah M, Azmi T, Hilmi M, Vidyadaran M, Bongso T, Nava Z, et al. Identification of crossbred buffalo genotypes and their chromosome segregation patterns. Genome. 1989;32(6):999–1002.

5. Deng T, Liang A, Liu J, Hua G, Ye T, Liu S, et al. Genome-Wide SNP Data Revealed the Extent of Linkage Disequilibrium, Persistence of Phase and Effective Population Size in Purebred and Crossbred Buffalo Populations. Frontiers in Genetics. 2019;9:688.

6. Yindee M, Vlamings B, Wajjwalku W, Techakumphu M, Lohachit C, Sirivaidyapong S, et al. Y-chromosomal variation confirms independent domestications of swamp and river buffalo. Animal genetics. 2010;41(4):433–5.

7. Lei C, Zhang W, Chen H, Lu F, Liu R, Yang X, et al. Independent maternal origin of Chinese swamp buffalo (Bubalus bubalis). Animal Genetics. 2007;38(2):97–102.

8. Kumar S, Nagarajan M, Sandhu JS, Kumar N, Behl V, Nishanth G. Mitochondrial DNA analyses of Indian water buffalo support a distinct genetic origin of river and swamp buffalo. Animal genetics. 2007;38(3):227–32. doi: 10.1111/j.1365-2052.2007.01602.x. PubMed PMID: 17459014.

9. Zhang Y, Lu Y, Yindee M, Li KY, Kuo HY, Ju YT, et al. Strong and stable geographic differentiation of swamp buffalo maternal and paternal lineages indicates domestication in the China/Indochina border region. Molecular ecology. 2016;25(7):1530–50.

10. Colli L, Milanesi M, Vajana E, Iamartino D, Bomba L, Puglisi F, et al. New insights on water buffalo genomic diversity and post-domestication migration routes from medium density SNP chip data. Frontiers in genetics. 2018;9:53.

11. Yue X-P, Li R, Xie W-M, Xu P, Chang T-C, Liu L, et al. Phylogeography and domestication of Chinese swamp buffalo. PloS one. 2013;8(2):e56552.

12. Wu FQ, Shen SK, Zhang XJ, Wang YH, Sun WB. Genetic diversity and population structure of an extremely endangered species: the world’s largest Rhododendron. AoB Plants. 2015;7:plu082.

13. Bertolini F, Schiavo G, Scotti E, Ribani A, Martelli PL, Casadio R, et al. High-throughput SNP discovery in the rabbit (Oryctolagus cuniculus) genome by next-generation semiconductor-based sequencing. Animal genetics. 2014;45(2):304–7. doi: 10.1111/age.12121. PubMed PMID: 24444082.

14. Blanca J, Esteras C, Ziarsolo P, Perez D, Ferna Ndez-Pedrosa V, Collado C, et al. Transcriptome sequencing for SNP discovery across Cucumis melo. BMC genomics. 2012;13:280. doi: 10.1186/1471-2164-13-280. PubMed PMID: 22726804; PubMed Central PMCID: PMC3473316.

15. CW W. SNP discovery in non-model organisms using 454 next generation sequencing. Methods Mol Biol. 2012;888(33-53).

16. Deng T, Pang C, Lu X, Zhu P, Duan A, Tan Z, et al. De Novo Transcriptome Assembly of the Chinese Swamp Buffalo by RNA Sequencing and SSR Marker Discovery. PloS one. 2016;11(1):e0147132. doi: 10.1371/journal.pone.0147132. PubMed PMID: 26766209; PubMed Central PMCID: PMC4713091.

17. Myakishev MV, Khripin Y, Hu S, Hamer DH. High-throughput SNP genotyping by allele-specific PCR with universal energy-transfer-labeled primers. Genome research. 2001;11(1):163–9. PubMed PMID: 11156625; PubMed Central PMCID: PMC311033.

18. Fan J-B, Chen X, Halushka MK, Berno A, Huang X, Ryder T, et al. Parallel genotyping of human SNPs using generic high-density oligonucleotide tag arrays. Genome Research. 2000;10(6):853–60.

19. Tosser-Klopp G, Bardou P, Bouchez O, Cabau C, Crooijmans R, Dong Y, et al. Design and characterization of a 52K SNP chip for goats. PloS one. 2014;9(1):e86227. doi: 10.1371/journal.pone.0086227. PubMed PMID: 24465974; PubMed Central PMCID: PMC3899236.

20. Porto-Neto LR, Sonstegard TS, Liu GE, Bickhart DM, Da Silva MV, Machado MA, et al. Genomic divergence of zebu and taurine cattle identified through high-density SNP genotyping. BMC genomics. 2013;14:876. doi: 10.1186/1471-2164-14-876. PubMed PMID: 24330634; PubMed Central PMCID: PMC4046821.

21. Manunza A, Zidi A, Yeghoyan S, Balteanu VA, Carsai TC, Scherbakov O, et al. A high throughput genotyping approach reveals distinctive autosomal genetic signatures for European and Near Eastern wild boar. PloS one. 2013;8(2):e55891. doi: 10.1371/journal.pone.0055891. PubMed PMID: 23460788; PubMed Central PMCID: PMC3584081.

22. Kujur A, Bajaj D, Upadhyaya HD, Das S, Ranjan R, Shree T, et al. A genomewide SNP scan accelerates trait-regulatory genomic loci identification in chickpea. Scientific reports. 2015;5:11166. doi: 10.1038/srep11166. PubMed PMID: 26058368; PubMed Central PMCID: PMC4461920.

23. Iamartino D, Nicolazzi EL, Van CT, Reecy JM, Fritzwaters ER, Koltes JE, et al. Design and validation of a 90K SNP genotyping assay for the water buffalo (Bubalus bubalis). PloS one. 2017;12(10):e0185220.

24. de Camargo GM, Aspilcueta-Borquis RR, Fortes MR, Porto-Neto R, Cardoso DF, Santos DJ, et al. Prospecting major genes in dairy buffaloes. BMC Genomics. 2015;16:872. doi: 10.1186/s12864-015-1986-2. PubMed PMID: 26510479; PubMed Central PMCID: PMC4625573.

25. Liu JJ, Liang AX, Campanile G, Plastow G, Zhang C, Wang Z, et al. Genomewide association studies to identify quantitative trait loci affecting milk production traits in water buffalo. Journal of dairy science. 2017.

26. El-Halawany N, Abdel-Shafy H, Shawky AEMA, Abdel-Latif MA, Al-Tohamy AFM, El-Moneim OMA, editors. Genome-wide association study for milk production in Egyptian buffalo. The International Symposium on Animal Functional Genomics; 2017.

27. Biswas S, Akey JM. Genomic insights into positive selection. TRENDS in Genetics. 2006;22(8):437–46.

28. Helyar SJ, Hemmer-Hansen J, Bekkevold D, Taylor M, Ogden R, Limborg M, et al. Application of SNPs for population genetics of nonmodel organisms: new opportunities and challenges. Molecular ecology resources. 2011;11:123–36.

29. Gouveia JJdS, Silva MVGBd, Paiva SR, Oliveira SMPd. Identification of selection signatures in livestock species. Genetics and molecular biology. 2014;37(2):330–42.

30. Qanbari S, Pimentel E, Tetens J, Thaller G, Lichtner P, Sharifi A, et al. A genome-wide scan for signatures of recent selection in Holstein cattle. Animal genetics. 2010;41(4):377–89.

31. Qanbari S, Gianola D, Hayes B, Schenkel F, Miller S, Moore S, et al. Application of site and haplotype-frequency based approaches for detecting selection signatures in cattle. BMC Genomics. 2011;12(1):318.

32. Barendse W, Harrison BE, Bunch RJ, Thomas MB, Turner LB. Genome wide signatures of positive selection: the comparison of independent samples and the identification of regions associated to traits. BMC Genomics. 2009;10(1):178.

33. Rubin C-J, Megens H-J, Barrio AM, Maqbool K, Sayyab S, Schwochow D, et al. Strong signatures of selection in the domestic pig genome. Proceedings of the National Academy of Sciences. 2012;109(48):19529–36.

34. Kijas JW, Lenstra JA, Hayes B, Boitard S, Neto LRP, San Cristobal M, et al. Genome-wide analysis of the world’s sheep breeds reveals high levels of historic mixture and strong recent selection. PLoS Biology. 2012;10(2):e1001258.

35. Mokhber M, Moradi-Shahrbabak M, Sadeghi M, Moradi-Shahrbabak H, Stella A, Nicolzzi E, et al. A genome-wide scan for signatures of selection in Azeri and Khuzestani buffalo breeds. BMC Genomics. 2018;19(1):449.

36. Fariello MI, Boitard S, Naya H, SanCristobal M, Servin B. Detecting signatures of selection through haplotype differentiation among hierarchically structured populations. Genetics. 2013;193(3):929–41.

37. Walugembe M, Bertolini F, Dematawewa CMB, Reis MP, Elbeltagy AR, Schmidt CJ, et al. Detection of selection signatures among Brazilian, Sri Lankan, and Egyptian chicken populations under different environmental conditions. Frontiers in Genetics. 2018;9:737.

38. Zhang Y SD, Yu Y, Zhang Y. Genetic variation and divergence among swamp buffalo, river buffalo and cattle: a microsatellite survey on five populations in China. Asian Austral J Anim Sci. 2008;21:1238–43.

39. Barker JS, Moore SS, Hetzel DJ, Evans D, Tan SG, Byrne K. Genetic diversity of Asian water buffalo (Bubalus bubalis): microsatellite variation and a comparison with protein-coding loci. Animal genetics. 1997;28(2):103–15. PubMed PMID: 9172308.

40. Zhang Y, Sun D, Yu Y, Zhang Y. Genetic diversity and differentiation of Chinese domestic buffalo based on 30 microsatellite markers. Animal genetics. 2007;38(6):569–75. doi: 10.1111/j.1365-2052.2007.01648.x. PubMed PMID: 17980000.

41. Zhang Y, Lu Y, Yindee M, Li KY, Kuo HY, Ju YT, et al. Strong and stable geographic differentiation of swamp buffalo maternal and paternal lineages indicates domestication in the China/Indochina border region. Molecular ecology. 2016;25(7):1530–50. doi: 10.1111/mec.13518. PubMed PMID: 26677084.

42. Kathiravan P, Kataria R, Mishra B, Dubey P, Sadana D, Joshi B. Population structure and phylogeography of Toda buffalo in Nilgiris throw light on possible origin of aboriginal Toda tribe of South India. Journal of Animal Breeding and Genetics. 2011;128(4):295–304.

43. Iamartino D, Nicolazzi EL, Van Tassell CP, Reecy JM, Fritz-Waters ER, Koltes JE, et al. Design and validation of a 90K SNP genotyping assay for the water buffalo (Bubalus bubalis). PLoS One. 2017;12(10):e0185220. doi: 10.1371/journal.pone.0185220. PubMed Central PMCID: PMC5628821.

44. Smitz N, Berthouly C, Cornelis D, Heller R, Van Hooft P, Chardonnet P, et al. Pan-African genetic structure in the African buffalo (Syncerus caffer): investigating intraspecific divergence. PloS one. 2013;8(2):e56235. doi: 10.1371/journal.pone.0056235. PubMed PMID: 23437100; PubMed Central PMCID: PMC3578844.

45. Zhang Y, Vankan D, Zhang Y, Barker JS. Genetic differentiation of water buffalo (Bubalus bubalis) populations in China, Nepal and south-east Asia: inferences on the region of domestication of the swamp buffalo. Animal genetics. 2011;42(4):366–77. doi: 10.1111/j.1365-2052.2010.02166.x. PubMed PMID:21749419.

46. Vijh RK, Tantia MS, Mishra B, Bharani Kumar ST. Genetic relationship and diversity analysis of Indian water buffalo (Bubalus bubalis). Journal of animal science. 2008;86(7):1495–502. doi: 10.2527/jas.2007-0321. PubMed PMID: 18344309.

47. Mishra B, Dubey P, Prakash B, Kathiravan P, Goyal S, Sadana D, et al. Genetic analysis of river, swamp and hybrid buffaloes of north-east India throw new light on phylogeography of water buffalo (Bubalus bubalis). Journal of Animal Breeding and Genetics. 2015;132(6):454–66.

48. Purfield DC, Berry DP, Sinead MP, Bradley DG. Runs of homozygosity and population history in cattle. BMC genetics. 2012;13(1):1–11.

49. Bjelland DW, Weigel KA, Vukasinovic N, Nkrumah JD. Evaluation of inbreeding depression in Holstein cattle using whole-genome SNP markers and alternative measures of genomic inbreeding. Journal of dairy science. 2013;96(7):4697–706.

50. Ferencakovic M, Sölkner J, Curik I. Estimating autozygosity from high-throughput information: effects of SNP density and genotyping errors. Genetics Selection Evolution. 2013;45(1):42. PubMed Central PMCID: PMC4176748.

51. Makina SO, Muchadeyi FC, van Marle-Koster E, Taylor JF, Makgahlela ML, Maiwashe A. Genome-wide scan for selection signatures in six cattle breeds in South Africa. Genetics, selection, evolution : GSE. 2015;47:92. doi: 10.1186/s12711-015-0173-x. PubMed PMID: 26612660; PubMed Central PMCID: PMC4662009.

52. Darshi M, Mendiola VL, Mackey MR, Murphy AN, Koller A, Perkins GA, et al. ChChd3, an inner mitochondrial membrane protein, is essential for maintaining crista integrity and mitochondrial function. Journal of Biological Chemistry. 2011;286(4):2918–32.

53. Atmuri V, Martin DC, Hemming R, Gutsol A, Byers S, Sahebjam S, et al. Hyaluronidase 3 (HYAL3) knockout mice do not display evidence of hyaluronan accumulation. Matrix Biology. 2008;27(8):653–60.

54. Badylak SF, Freytes DO, Gilbert TW. Extracellular matrix as a biological scaffold material: structure and function. Acta biomaterialia. 2009;5(1):1–13.

55. Theocharis AD, Skandalis SS, Gialeli C, Karamanos NK. Extracellular matrix structure. Advanced Drug Delivery Reviews. 2016;97:4–27.

56. Asnaghi L, White DT, Key N, Choi J, Mahale A, Alkatan H, et al. ACVR1C/SMAD2 signaling promotes invasion and growth in retinoblastoma. Oncogene. 2019;38(12):2056–75.

57. Saxton RA, Sabatini DM. mTOR signaling in growth, metabolism, and disease. Cell. 2017;169(2):361–71.

58. Hu Y, Su H, Liu C, Wang Z, Huang L, Wang Q, et al. DEPTOR is a direct NOTCH1 target that promotes cell proliferation and survival in T-cell leukemia. Oncogene. 2017;36(8):1038–47.

59. Tőkési N, Lehotzky A, Horváth I, Szabó B, Oláh J, Lau P, et al. TPPP/p25 promotes tubulin acetylation by inhibiting histone deacetylase 6. Journal of Biological Chemistry. 2010;285(23):17896–906.

60. Zhou W, Wang X, Li L, Feng X, Yang Z, Zhang W, et al. Depletion of tubulin polymerization promoting protein family member 3 suppresses HeLa cell proliferation. Molecular Cellular Biochemistry. 2010;333(1-2):91–8.

61. Nishimura S, Watanabe T, Mizoshita K, Tatsuda K, Fujita T, Watanabe N, et al. Genome-wide association study identified three major QTL for carcass weight including the PLAG1-CHCHD7 QTN for stature in Japanese Black cattle. BMC Genetics. 2012;13(1):40.

62. Li Y, Yang K, Yang W, Chu L, Chen C, Zhao B, et al. Identification of QTL and Qualitative Trait Loci for Agronomic Traits Using SNP Markers in the Adzuki Bean. Frontiers in Plant Science. 2017;8:840.

63. Liu J, Liang A, Campanile G, Plastow G, Zhang C, Wang Z, et al. Genome-wide association studies to identify quantitative trait loci affecting milk production traits in water buffalo. Journal of Dairy Science. 2018;101(1):433–44.

64. Tu C-F, Wu M-Y, Lin Y-C, Kannagi R, Yang R-B. FUT8 promotes breast cancer cell invasiveness by remodeling TGF-β receptor core fucosylation. Breast Cancer Research. 2017;19(1):111.

65. Fujii-Kuriyama Y, Kawajiri K. Molecular mechanisms of the physiological functions of the aryl hydrocarbon (dioxin) receptor, a multifunctional regulator that senses and responds to environmental stimuli. Proc Jpn Acad Ser B Phys Biol Sci. 2010;86(1):40–53.

66. Oshimori N, Li X, Ohsugi M, Yamamoto T. Cep72 regulates the localization of key centrosomal proteins and proper bipolar spindle formation. EMBO Journal. 2009;28(14):2066–76.

67. Vincze O, Tökési N, Oláh J, Hlavanda E, Zotter Á, Horváth I, et al. Tubulin polymerization promoting proteins (TPPPs): members of a new family with distinct structures and functions. Biochemistry. 2006;45(46):13818–26.

68. Basham KJ, Leonard CJ, Kieffer C, Shelton DN, McDowell ME, Bhonde VR, et al. Dioxin exposure blocks lactation through a direct effect on mammary epithelial cells mediated by the aryl hydrocarbon receptor repressor. Toxicological Sciences. 2014;143(1):36–45.

69. Bionaz M, Loor JJ. Gene networks driving bovine mammary protein synthesis during the lactation cycle. Bioinformatics and biology insights. 2011;5:BBI. S7003.

70. Chang CC, Chow CC, Tellier LC, Vattikuti S, Purcell SM, Lee JJ. Second-generation PLINK: rising to the challenge of larger and richer datasets. GigaScience. 2015;4:7. doi: 10.1186/s13742-015-0047-8. PubMed PMID: 25722852; PubMed Central PMCID: PMC4342193.

71. Patterson N, Price AL, Reich D. Patterson N, Price AL, Reich D. Population structure and eigenanalysis. PLoS Genet 2: 2074–2093. Plos Genetics. 2007;2(12):e190.

72. Cummings MP. PHYLIP (Phylogeny Inference Package): John Wiley & Sons, Inc.; 2004. 164–6 p.

73. Tamura K, Stecher G, Peterson D, Filipski A, Kumar S. MEGA6: Molecular Evolutionary Genetics Analysis version 6.0. Molecular biology and evolution. 2013;30(12):2725–9. Epub 2013/10/18. doi: 10.1093/molbev/mst197. PubMed PMID: 24132122; PubMed Central PMCID: PMCPMC3840312.

74. Alexander DH, Novembre J, Lange K. Fast model-based estimation of ancestry in unrelated individuals. Genome research. 2009;19(9):1655–64.

75. Excoffier L, Lischer HE. Arlequin suite ver 3.5: a new series of programs to perform population genetics analyses under Linux and Windows. Molecular ecology resources. 2010;10(3):564–7. doi: 10.1111/j.1755-0998.2010.02847.x. PubMed PMID: 21565059.

76. Jombart T. adegenet: a R package for the multivariate analysis of genetic markers. Bioinformatics. 2008;24(11):1403–5. doi: 10.1093/bioinformatics/btn129. PubMed PMID: 18397895.

77. Zhang C, Dong S-S, Xu J-Y, He W-M, Yang T-L. PopLDdecay: a fast and effective tool for linkage disequilibrium decay analysis based on variant call format files. Bioinformatics. 2018;35(10):1786–8.

78. Biscarini F, Cozzi P, Gaspa G, Marras G. detectRUNS: Detect Runs of Homozygosity and Runs of Heterozygosity in Diploid Genomes. 2018.

79. Fariello MI, Boitard S, Naya H, Sancristobal M, Servin B. Detecting signatures of selection through haplotype differentiation among hierarchically structured populations. Genetics. 2013;193(3):929–U448.

80. Coreteam R. R: A language and environment for statistical computing. Computing. 2018;1:12–21.

